# Preserving Integrity: Innovative *In Vitro* Methods for Extracellular Matrix Decellularization and Collagen Purification

**DOI:** 10.1101/2025.08.22.671770

**Authors:** Uliana Bashtanova, Rui Li, Ieva Goldberga, Kathryn Gerl, Kristen Paige Burgess, Annika Janine Wegner-Repke, Melinda Jane Duer

**Affiliations:** Yusuf Hamied Department of Chemistry, University of Cambridge, Lensfield Rd, Cambridge CB2 1EW, UK

**Keywords:** Collagen, triple helix, extracellular matrix, decellularization, cytoskeleton, vinblastin, latrunculin, orthovanadate, biomaterials

## Abstract

**Background:** In tissue engineering and cell therapy development, synthetic biomaterials are frequently supplemented with collagen or other extracellular matrix (ECM) components to enhance biocompatibility. To support these applications, novel methods for collagen purification and ECM decellularization were developed, with a focus on preserving the structural and biochemical integrity of the final products.

**Results:** The effectiveness of these methods was validated using solid-state NMR and fluorescence spectroscopy, bright-field and confocal microscopy, amino acid analysis, and transmission electron microscopy. Intact cells were dislodged from ECM-producing cultures through the application of cytoskeleton-targeting drugs, while the native protein composition of the ECM was maintained. In parallel, collagen purified using chymotrypsin was shown to retain its native triple-helical structure and post-translational modifications.

**Conclusions:** Both techniques are broadly applicable to various cell types capable of producing collagen and/or ECM *in vitro*, thereby expanding the availability of species- and tissue-specific sources. These advances hold particular promise for human-relevant tissue engineering and drug discovery applications.

## INTRODUCTION

The extracellular matrix (ECM) is the non-cellular component present in all tissues and organs, providing essential physical scaffolding for cells. The main structural components of the ECM are fibrillar collagen proteins, triple helical molecules that organize into ordered arrays forming fibrils 10s – 100s of nanometres in diameter, providing stiffness and regulating cell adhesion or migration [1,2]. The fibrous ECM network is further supported by other fibril and filament-forming proteins, including elastin, fibrillin, fibronectin, and laminin. These protein networks are coupled together by linker ECM proteins including tenascin, non-fibrous collagens and certain types of proteoglycans. The associations among these proteins not only maintain the 3D architecture of the ECM, as well as the orientation of collagen and elastin fibres, but also preserve the ECM’s elasticity, tensile strength, and porosity required by each tissue.

Equally important, they regulate ligand presentation for cell surface integrins, growth factors, and signalling molecules. Proteoglycans, together with hyaluronic acid, form a hydrated gel that provides pH buffering, tissue hydration, reservoirs of cations and chemokines, and lubrication [3,4]. The types of ECM proteins expressed, along with their folding, crosslinking, and post-translational modifications, are species-, organ-, tissue-, disease- and age-specific [5]. The complexity of these protein combinations, their post-translational modifications, and their characteristic 3D architecture within the ECM is difficult to replicate using fabrication methods. As a result, seeding cells onto protein beds derived from native ECM has become increasingly desirable over the past decade [6].

The first step in the ECM preparation process is a removal of cellular material. Numerous decellularization methods have been developed, which can be divided into physical, chemical, and enzymatic approaches [6]. Physical methods include freeze-thaw cycling and/or high hydrostatic pressure. They preserve structure of ECM, but on their own, are ineffective for complete decellularization and typically combined with chemical and enzymatic approaches [7]. Chemical reagents include acids and bases, hypo- or hypertonic solutions, and detergents. They facilitate the solubilization of cellular membranes and dissociation of DNA, which otherwise bind to the ECM after cell lysis, and the disruption of lipid-lipid/protein interactions [8]. Proteolytic enzymes are also used to ensure the removal of specific cellular protein or genetic material and sample enrichment of the ECM protein of interest [6].

Recovering the complete ECM—i.e., retaining its key proteins, glycans, and lipids that influence cell physiology—is enormously challenging and is difficult to achieve with current physical and chemical approaches. This is partly because these methods rely on cell lysis via physical disruption and/or chemical solubilization of cellular membranes, which leads to contamination of the ECM with cellular lipids and DNA, as well as the probable loss of soluble ECM proteins such as proteoglycans [9] and tenascins. We have previously demonstrated that conventional approaches, including freeze-thaw cell lysis, detergent treatment, and DNase digestion, are insufficient for the complete removal of DNA and nuclear membranes from the ECM [10]. The extraction of collagen alone—as a major scaffolding and anchoring protein—is also neither straightforward nor chemically mild process. It is typically achieved either by solubilization in acid (for tissues where immature aldimine crosslinks predominate) or by pepsin proteolysis of non-triple-helical domains (for tissues with predominantly mature crosslinks) [1,2] and then reconstituted. Clearly, reconstituted collagen, which was initially denatured by acid or digested by pepsin, cannot be considered native because pepsin proteolysis destroys the initial fibril structure, and acid hydrolysis breaks glycosidic bonds introduced during post-translational modifications [1,2].

In this paper we report a new method of collagen purification from *in vitro*-grown ECM. Our collagen clean-up method uses chymotrypsin to digest non-collagenous proteins, while preserving the collagen fibril structure and likely maintaining glycosidic bonds [11]. Such collagen can be further processed into gels, sponges, films, fibres, and scaffolds or combined with synthetic biomaterials to tailor their mechanical and biological properties.

Beyond this, we propose a completely novel approach to ECM decellularization. Our method employs cytoskeleton-targeting drugs to induce loss of adhesion in living cells, followed by their dislocation from the ECM—leveraging the cells’ physiological capacity for detachment. This approach delivers the most structurally and biochemically ‘unspoiled’ native ECM reported to date. We anticipate that the preserved ECM will be highly valuable for studies requiring a native matrix [7]. Additionally, it may serve as a reference material for the development of synthetic ECM-like biomaterials, or be combined with synthetic polymers in applications where 3D ECM scaffolds are essential [6,7,12,13].

## MATERIALS AND METHODS

### Cell Cultures

The bovine aorta vascular smooth muscle cells (**BVSMC**) were obtained from Prof. Catherine Shanahan, King’s College London. Passages #4-8 were used for the experiments. Cell culture flasks were incubated at 37°C in a humidified atmosphere with 95% air and 5% CO2, and the culture medium was refreshed every 2-3 days. Dulbecco’s Modified Eagle Medium (DMEM; Gibco) with 4.5 g/L glucose was supplemented with 10% foetal calf serum (Pan-Biotech) and 1% L-Penicillin-Streptomycin-Glutamine solution (Gibco). A stock solution of L-ascorbic acid (Sigma-Aldrich) was prepared in water, aliquoted, and stored frozen at −20°C for no longer than 2 months. It was added to the flask at a final concentration of 50 μg/mL with each media change, after the cells reached 100% confluency. ECM was harvested when the cells produced a dense matrix that began to peel off the surface of the tissue culture flask, typically around 25 days after cell seeding.

The parent cells of the foetal sheep osteoblasts (**FSOB**) were obtained from Dr Rakesh Rajan, Department of Chemistry, University of Cambridge and grown the same way as BVSMC.

Mouse preosteoblasts (**MC3T3-E1**) were acquired from ATCC and cultured in Minimum Essential Medium (α-MEM; Thermo Fisher) supplemented with 10% foetal calf serum and 1% L-Penicillin-Streptomycin-Glutamine solution (Gibco). Once the cells reached confluency, 50ug/ml L-ascorbic acid in Dulbecco’s phosphate buffered saline (DPBS; Thermo Fisher) was added to the growth media for 19 days. The ECM was harvested after a total of 5 weeks of growth.

The mouse pancreatic stellate cells (**PSC4**) were obtained from Dr Giulia Biffi, CRUK [14]. DMEM with 4.5 g/L glucose was supplemented with 5% foetal bovine serum and 1% PSG. As above, the media was supplemented with 50 µg/mL ascorbic acid throughout the 3-week treatment post-confluence.

### Freeze-thaw Cell Lysis

The flask with cells was placed in a freezer at −80°C overnight, and the cells were lysed by thawing the flask at room temperature for 30 minutes. The cell debris was then removed by repeated washes with PBS. The collected ECM in PBS was transferred to a Falcon tube. Samples were stored at −20°C prior to any further procedures.

### Detergent Cell Lysis

Medium was aspirated and cell washed twice with PBS. 10 mL of detergent solution [15] (0.5% Triton X-100, 20 mM NH₄OH in PBS) was added to a flask with cell cultures and incubated at 37°C for 10 minutes. ECM was dislodged by gently swirling the flask in the presence of detergent, transferred to a 50 mL Falcon tube, and supplemented with 10 mL of PBS. The tube was then left at 4°C overnight. To remove detergent and cell debris, the ECM was washed with PBS three times and then incubated with DNase (Thermo Fisher Scientific, 10 μg/mL, 1:1000 dilution in its reaction buffer) at 37°C for 30 minutes. DNase was washed out with PBS before transferring the ECM to a fresh Falcon tube. Samples were stored at −20°C prior to any further procedures.

### Collagen Purification

To remove non-collagenous proteins and extract collagen, a flask of ECM, partially decellularized either by freeze-thaw lysis or detergent lysis, was incubated with chymotrypsin (0.1 mg, Sigma) in 2 mL of buffer (100 mM Tris-HCl, 10 mM CaCl₂, pH 7.8 in water) at 37°C for different time intervals (ranging from 5 minutes to 24 hours). The ECM was then thoroughly washed with PBS. To remove digested proteins, the sample was incubated with a Tween-20 solution (0.25% in 2 mL PBS) at 37°C overnight and thoroughly washed again. Tween-20 was an essential step in BVSMC collagen cleanup, but not in FSOB. The sample was stored at −20°C in a Falcon tube until further processing.

### ECM Decellularization by Cytoskeleton-targeting Drugs

It was important to keep the cells alive during the decellularization procedure (see below for the description of different treatments). Therefore, during the procedure, the growth medium supplemented with glucose and FBS was used, and the flasks were kept in the cell incubator. To remove as many rounded cells as possible, the flask was firmly tapped for 5 minutes against the palm of the hand, but care was taken not to tear the ECM apart.

#### Vinblastine treatment

Vinblastine sulphate (Fisher Scientific; Bio-Techne) was dissolved in Milli-Q water at room temperature and then filter sterilized. The 12 mM stock solution was diluted to final concentrations by directly adding it to the flasks with cells. The vinblastine-containing medium was refreshed daily after the rounded cells were shaken off and counted. Treatment typically continued for 3-4 days until no floating cells were observed under the microscope. The stock solution of vinblastine was stored in the fridge for 2-3 weeks.

#### Sodium orthovanadate (Na3VO4) & vinblastine treatment

Na₃VO₄ (Fisher Scientific) was solubilized in cell growth medium preheated to 37°C. The maximal solubility could be different in different media: it was found to be 35 mM in α-MEM and 50 mM in DMEM growth medium. Full solubilization took 30 to 60 minutes at 37°C. The Na₃VO₄-containing medium was added to flasks in place of the usual medium (35 mM), then spiked with vinblastine to achieve a final concentration of 360 µM for BVSMC, 80 or 250 µM for PSC. The vinblastine stock solution was prepared as described above. The treatment usually started before midday and lasted overnight. The following morning, rounded cells were shaken off and counted, and the drug-containing medium was replaced with normal medium. After 3-4 hours, cells were shaken off again, counted, and then treated for a second overnight time period. Typically, 3 overnight treatments were sufficient to dislodge all cells from the ECM.

#### Latrunculin B & Na3VO4 & vinblastine sulphate treatment

Vinblastine sulphate and Na3VO4 were prepared as previously described. Latrunculin B (Merck) stock solution was prepared by dissolving the compound in DMSO to a concentration of 20ug/ul and stored at −20°C. On the first day of treatment, cells were incubated with 120 µM vinblastine sulphate in α-MEM for 1 hour. Latrunculin B was then added directly to the vinblastine-treated medium to a final concentration of 62 µM, and incubation continued for an additional 1.5 hours. The drug-containing media was removed and replaced with 35 mM Na3VO4 in α-MEM for 2.5 hours. Finally, the Na3VO4-treated media was removed and replaced with pure α-MEM overnight. The following morning, rounded cells were gently shaken off and counted, taking care to avoid vigorous agitation to prevent detachment of the matrix from the flask. From this point forward, the flask was gently shaken, and the number of lifted cells were counted between each medium change unless otherwise specified. The media was then replaced with α-MEM containing 120 µM vinblastine sulphate and 62 µM Latrunculin B for 2.5 hours. This was followed by replacement with 35mM Na3VO4 in α-MEM for 3 hours, after which it was replaced with fresh α-MEM overnight. On the final day of treatment, the medium was replaced with α-MEM containing 120 µM vinblastine sulphate and 62 µM Latrunculin B for a final 2.5-hour incubation. The decellularization procedure was completed by removing the drug-treated medium and rinsing the samples twice with PBS pH 7.4.

### Amino Acid Analysis

Amino acid analysis was provided by the Department of Biochemistry, University of Cambridge. Dried samples were provided to the service for analysis. The procedure is described in [16] with slight modifications for handling the extracellular matrix samples. Briefly, to carefully weighted, lipolyzed sample a solution containing L-norleucine (100 μL, 250 nM in 0.1 HCl, Sigma) was added as an internal standard.. The mixture was incubated for 1 h at 4 °C and centrifuged at 3000 g for 15 min. The sample was transferred to a pyrolysed tube, where the residual liquid is removed via centrifugal evaporator (SpeedVacc). Then the sample was placed in a hydrolysis vial containing a mixture of concentrated HCl and phenol (0.5 mL) together with dodecanthiol (0.68 μL, Sigma) at 4 °C. The hydrolysis vial was evacuated and flushed with argon four times. Finally, the vial was placed in an oven at 115 °C for 22 h for amino acid hydrolysis. The sample was placed in a desiccator over solid NaOH for 40 min until the acid was removed. The sample was then dissolved in sodium citrate loading buffer (pH 2.2, Sigma), centrifuged at 3000 g for 15 min and then filtered through a 0.2-μm filter. The filtrate was injected into a loading capsule placed in a Pharmacia Alpha Plus series amino acid analyser (Biochrom Ltd, Cambridge, UK). Chromatography was performed on a sodium system ion exchange resin eluting with buffers over the pH range 3.2 to 5.45. Peak detection was achieved by mixing eluate with ninhydrin at 135 °C and measuring the absorbance at 570 and 440 nm. The area of the signals was taken of each amino acid and compared with respect to the internal standard and scaled against it to find mol%.

### Fluorescence Spectroscopy

Measurements were performed using a Cary Eclipse Fluorescence Spectrophotometer (Agilent, Santa Clara, CA, USA) with excitation at 275 nm and at 295 K. ECM was gently placed in a micro quartz cuvette with a path length of 10 mm and a chamber volume of 700 μL (SUPRASIL®, Hellma Analytics, Mülheim, Germany). The water solvent background was recorded and subtracted from the ECM spectra.

### Transmission Electron Microscopy (TEM) Imaging

ECM samples are gentled sonicated to better disperse sample in solution. 5 μl of solution was adsorbed onto glow-discharged 400 mesh copper/carbon-film grids (EM Resolutions) for about 2 min. Grids were rinsed on two drops of DIW and negative staining was performed using a 2 % aqueous uranyl acetate solution. Grids were viewed in a FEI Tecnai G^2^ electron microscope run at 200 keV using a 10 μm objective aperture. Images were acquired using AMT Camera software.

### Sample Preparation for ssNMR

Collagen samples were freeze-dried for 72 hours, yielding a white powder that could be easily packed into 4 mm zirconia MAS rotors (Bruker). As a reference pure collagen type I from bovine Achilles tendon (Sigma) was used.

However, freeze-dry conditions may compromise the structural integrity of decellularized ECM. Therefore, a gentler sample preparation method was used for our decellularized ECM samples, which involved freezing them in a hydrated state, as detailed below.

The non-decellularized ECM was attached to the bottom of the culture flask and was carefully stripped, starting from the edges and progressing toward the centre. In most cases, the ECM could be lifted in one piece. Minimal disruption was critical to avoid biochemical alterations caused by cellular proteases. Following decellularization, the ECM became free-floating in the medium.

Both non-decellularized and decellularized ECM samples were washed twice in serum- and glucose-free medium, followed by deionized (DI) water to remove residual salts, which can suppress radiofrequency (rf) currents during NMR. Excess liquid was removed using lint-free paper. Samples were then packed into a 3.2 mm zirconia Bruker MAS rotors with Vespel caps, frozen on dry ice, and stored at −80 °C. All steps were performed with care to prevent thawing and potential protein unfolding.

For NMR analysis of dislodged cells, samples were collected after the first day of treatment, centrifuged at 1400 rpm for 5 minutes, washed with medium lacking serum and glucose, centrifuged again, and packed into rotors. They were then frozen on dry ice and stored at −80 °C. As with ECM samples, every precaution was taken to prevent defrosting.

### Sample Staining for Confocal Microscopy

The control, non-decellularised sample, which was attached to the bottom of the flask, was carefully detached using a cell scraper as described in the previous section. The decellularised sample was already suspended in the culture medium and did not require scraping. Both samples, now floating in cell culture medium, were transferred to separate 100mm^2^ petri dishes, and rinsed once with 1X PBS (pH 7.4). The matrix samples were then suspended in fresh PBS (pH 7.4), and tweezers were used to gently spread them out, removing any folds or creases. Once the matrices were evenly spread with minimal wrinkles, a glass microscope slide was inserted into each petri dish. The dishes were carefully tilted to allow the matrix to evenly spread across the surface of the slide. Next, a working solution of 10 µM Hoechst 33342 (Thermo Fisher Scientific) was prepared in 1X PBS (pH 7.4). The microscope slides, with matrices on top, were briefly removed the dishes, and the remaining PBS was discarded. The microscope slides were returned to the edges of the petri dishes, and the Hoechst solution was carefully added until the matrices were fully submerged. To ensure complete submersion, a small support was placed under one corner of each dish to slightly tilt it. The samples were incubated in Hoechst for 10 minutes in the dark. Following incubation, the slides were removed, and the matrices were gently adjusted to remove any remaining wrinkles. Finally, a large rectangular coverslip was placed over each sample, taking care to avoid introducing air bubbles. The samples were imaged using 405 nm excitation with a STELLARIS 5 inverted fluorescent microscope (Leica microsystems)

### ssNMR

All NMR data was acquired on a Bruker 9.4 T wide bore NMR magnet system with a ^1^H Larmor frequency of 400 MHz, which was equipped with an Avance I or Neo console. For the NMR spectra of commercial collagen and isolated collagen a 4 mm HX double resonance MASDVT probe (Bruker) was utilised. Whereas, NMR experiments on the ECM and detached cells were conducted with a 3.2 mm HCP triple resonance MASDVT Efree probe whose coil design is known to minimise sample heating of salt-rich samples. The ^13^C chemical shifts were indirectly referenced to the trimethylsilane scale by using α-glycine as external standard [δ(^13^Cα)=43.1 ppm]. Probe sensor temperatures were corrected for MAS-induced frictional heating by measuring the temperature-dependent T1(^79^Br) relaxation time to determine the actual sample temperature [17]. All temperatures stated in this study refer to the calibrated sample temperatures at the respective MAS rates.

^1^H-^13^C CP/MAS experiments were employed to enhance the sensitivity of the natural abundance samples by transferring polarisation from high-abundance ^1^H spins to the low-abundance, low gyromagnetic ratio ^13^C nuclei *via* dipolar coupling. A ^1^H 90° pulse of 2.1-2.5 µs was used, followed by a ramped contact pulse of 2.0 or 2.5 ms at radiofrequency field strengths of ∼90 kHz for ^1^H and 50-70 kHz for ^13^C, optimised to fulfil the Hartmann-Hahn matching condition. During acquisition SPINAL-64 ^1^H decoupling was applied at 90 kHz [18]. Recycle delays of 2-5 s were utilised depending on the sample.

NMR spectra were processed in TopSpin 4.5.0.

## RESULTS

### Collagen purification

Extracting collagen from a cell culture offers the advantage of full control over the animal species, cell type, growth conditions, and medium composition, so the resulting collagen exhibits the specific structural and biochemical characteristics, including post-translational modifications, relevant to the tissue which are impossible to achieve with the very limited selection of commercially-available collagen preparations. However, *in vitro* ECM contains many proteins other than collagen, and their removal is challenging, because commonly used proteolytic enzymes such as trypsin, papain, and pepsin can digest collagen along with the target proteins [19].

Thus, our first step was to identify an enzyme capable of removing non-collagenous proteins, including cellular proteins, from *in vitro* cell cultures with minimal disruption to collagen fibrils. Chymotrypsin was eventually chosen because it preferentially cleaves at the carbonyl side of amide bonds in large hydrophobic or aromatic amino acids, such as phenylalanine, tyrosine, tryptophan, and leucine, unless it is protected by neighbouring proline [20]. Collagen contains far fewer aromatic amino acids compared to other ECM proteins, and of those few, many have a neighbouring proline or hydroxyproline residue, resulting in fewer chymotrypsin cleavage sites than in other proteins. For example, in bovine collagen type I, only 2.3% of the amide bonds are theoretically cleavable, while in bovine fibronectin isoform 1, this number increases to 12.6% (Table 1). An additional advantage is that most enzymes do not readily cleave intact triple helical collagen, as the steric hindrance provided by the triple helix protects the proteolytic cleavage sites [1] and for effective proteolysis, collagen must first be denatured—through heat treatment or acid hydrolysis—to separate the collagen α chains in the triple helix [1]. Chymotrypsin activity is maximal at pH 7.8–8 and remains effective at the physiological pH of 7.4 [21]. The use of this gentle pH condition minimized the risk of collagen denaturation and reduced the exposure of cleavage sites to chymotrypsin.

**Table 1.**
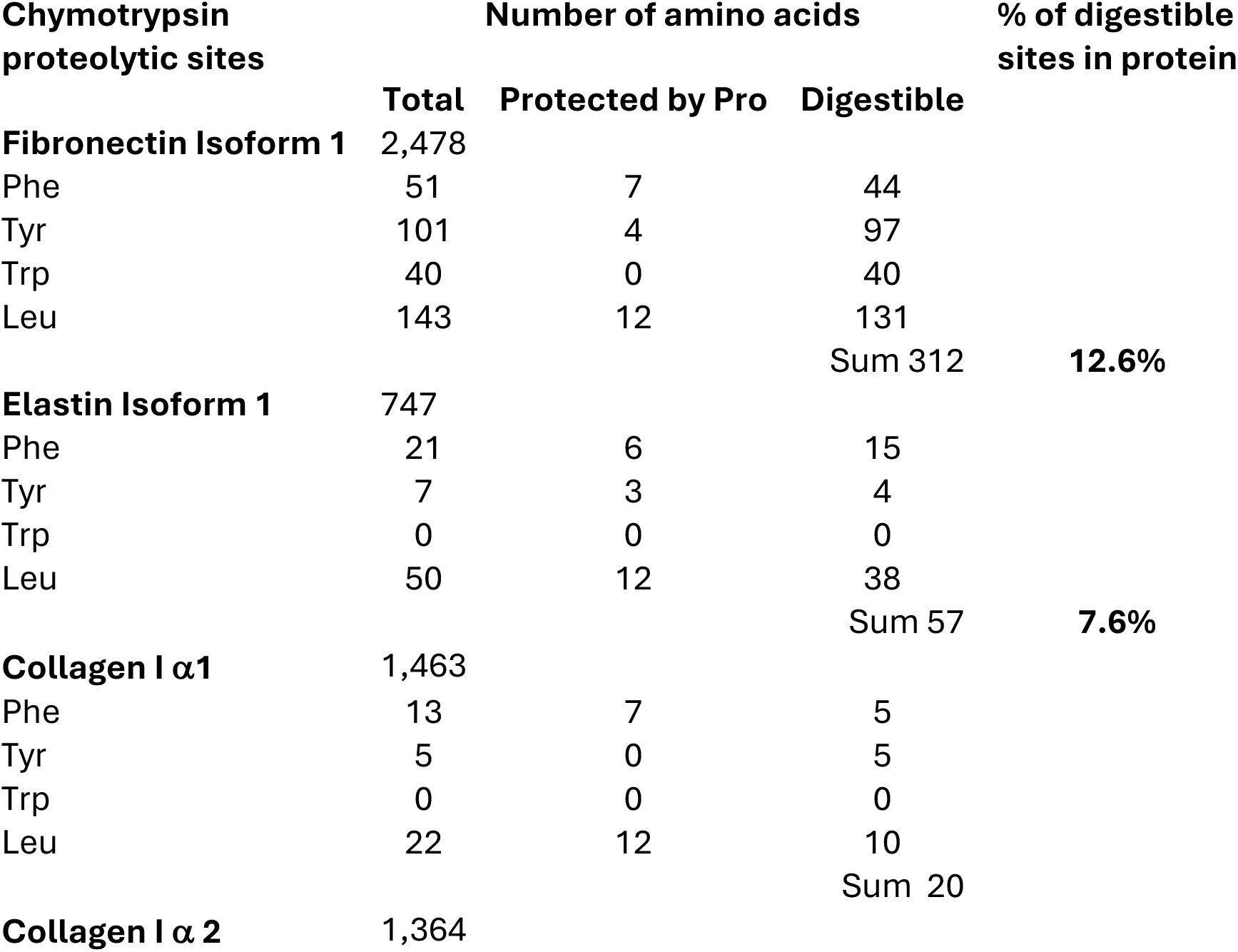
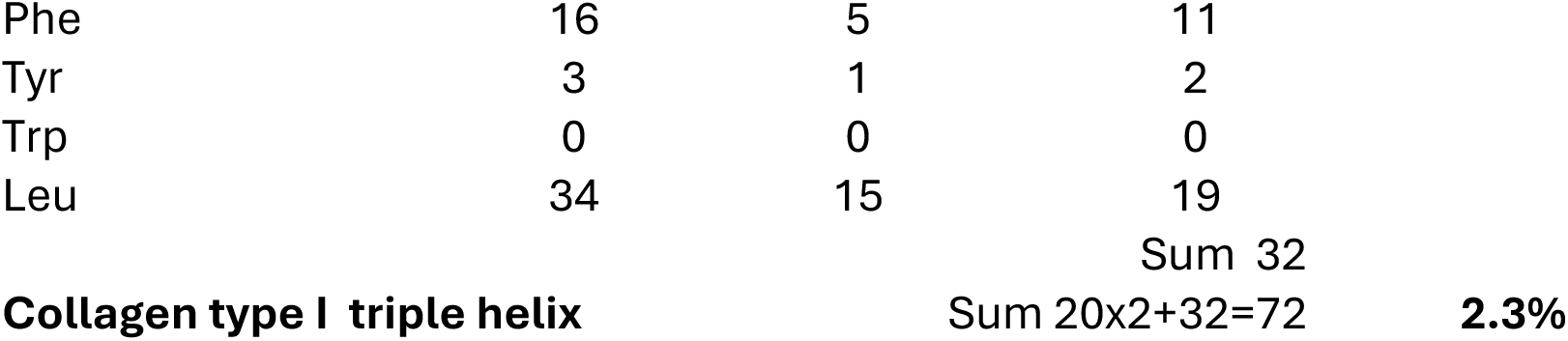
Chymotrypsin-cleavable sites in selected fibrillar bovine ECM proteins.

For demonstration purposes, the calculation assumes that chymotrypsin cleaves only at four specific amino acids. Residue counts were determined from sequences published in the UniProt protein sequence database [22]. Identifiers for Bovine Collagen Type I were P02453 (α1) and P02465 (α2); for Bovine Fibronectin Isoform 1 – P07589-1; and for Bovine Elastin Isoform 1 – P04985-1.

For the specific purpose of collagen purification, perfect decellularization was not essential, as the nucleus and membrane proteins were obviously sensitive to chymotrypsin digestion, along with non-collagenous ECM proteins. Therefore, for collagen purification, we chose to use a conventional physical/chemical decellularization method – freeze-thaw cell lysis followed by detergent washing (see Methods), as it is quick and requires minimal resources.

Collagen autofluorescence was used to determine whether chymotrypsin could selectively remove non-collagenous proteins from ECM-producing cell cultures and to identify the optimal clean-up conditions. There are three fluorescent amino acids in proteins: tyrosine (Tyr), tryptophan (Trp) and phenylalanine (Phe). Tryptophan is absent in collagen, and phenylalanine has a low quantum yield, therefore its fluorescence is not detectable in the presence of tyrosine or tryptophan [23–25]. Thus, collagen autofluorescence in the UV-A region originates from excitation and emission from its tyrosine residues which are predominantly located within the collagen telopeptides [26]. Tyrosine and tryptophan differ substantially in their emission wavelengths: tyrosine has an excitation/emission maximum at 275/303 nm at neutral pH, while tryptophan has an excitation/emission maximum at 280/330–355 nm (depending on the polarity of the environment [23–25], resulting in a difference in the emission wavelength of at least 25 nm. Thus, monitoring tryptophan autofluorescence enabled us to track the chymotrypsin digestion of non-collagenous proteins, while monitoring tyrosine autofluorescence allowed us to stop the reaction if collagen began to be digested; the non-triple helical collagen telopeptides where most of the collagen tyrosine residues reside are the most susceptible part of the collagen molecule to chymotrypsin cleavage, so we expected tyrosine fluorescence to be a sensitive monitor of any unintended collagen digestion.

As expected, neither freeze-thaw cell lysis nor subsequent detergent washes of the ECM resulted in the removal of tryptophan autofluorescence, which had a very broad peak with maximum around 330 nm (Fig.1A). Different chymotrypsin incubation times were tested, and for BVSMC and FSOB ECM overnight incubation was found to be optimal for the selective digestion of non-collagenous proteins. After chymotrypsin digestion, tryptophan autofluorescence was no longer observed in the ECM, while tyrosine autofluorescence remained (Fig.1A, B), indicating the presence of collagen fibrils with preserved telopeptides. In the case of BVSMC ECM, the emission maximum of tyrosine autofluorescence was sharp at 295 nm (Fig.1B), whereas in the case of FSOB ECM, the fluorescence peak was broader with the maximum ranging from 300 to 310 nm (Fig.1A). This likely reflected differences in the ratio of the collagen types and/or their environments in the two cell cultures.

**Fig. 1.**
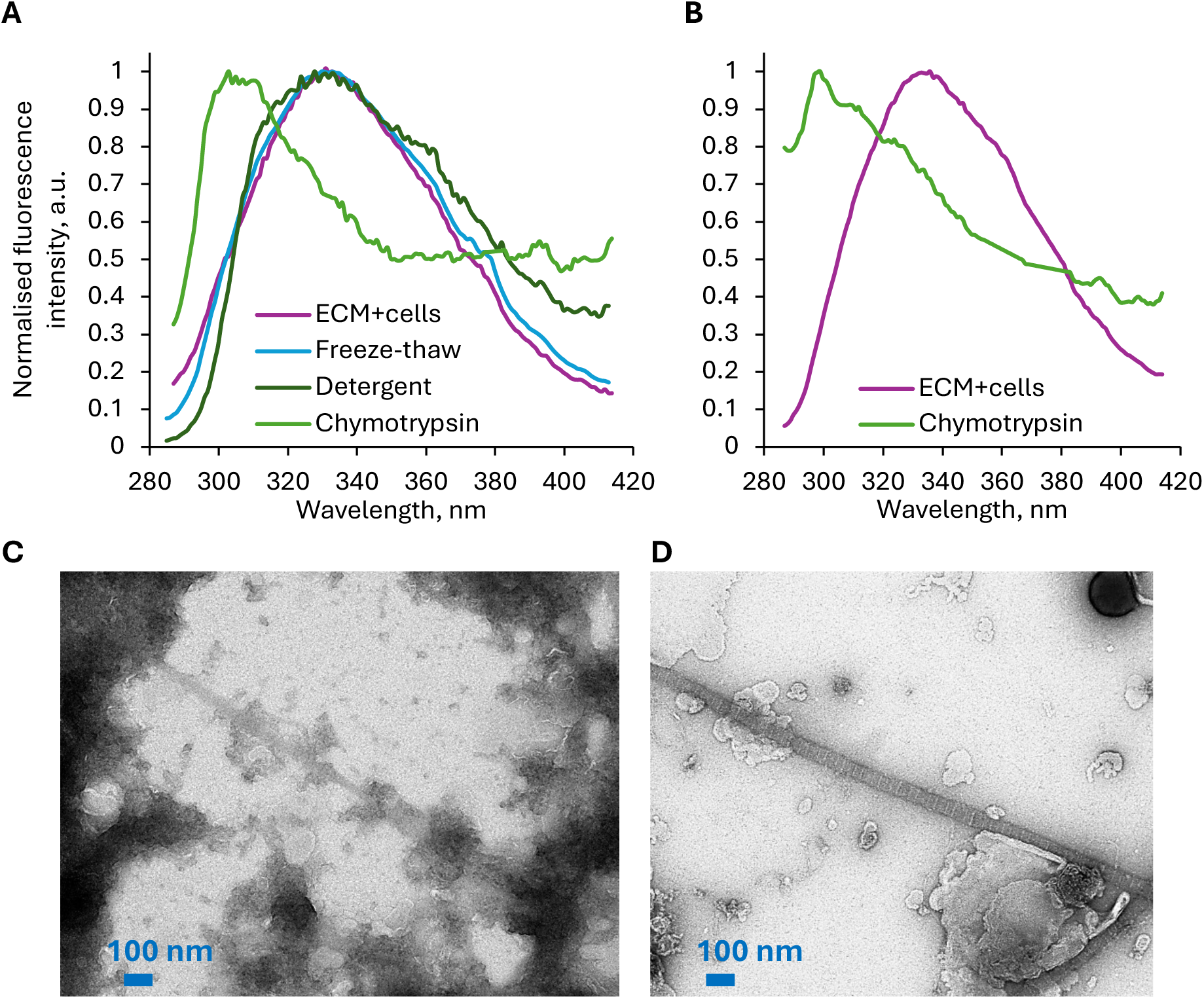
Collagen Isolation from *In Vitro*-Grown ECM. The effectiveness of collagen extraction was assessed by autofluorescence emission spectra (excitation at 275 nm) or transmission electron microscopy (TEM). The extraction steps included either freeze-thaw cell lysis (“freeze-thaw”), or Triton X-100 cell lysis (“detergent”), followed by chymotrypsin digestion (“chymotrypsin”). The control spectrum was represented by the cellular sheet before decellularization (“ECM+cells”). **A-B.** Tyrosine fluorescence (emission maximum at 300 nm), specific for collagen, was only detected after the chymotrypsin step in FSOB (**A**) and BVSMC (**B**). **C-D.** TEM images of BVSMC ECM demonstrate that after the freeze-thaw or detergent steps, fibrils, possibly collagen, were visible, but their banding pattern was obscured by numerous non-fibrillar proteins (**C**). After chymotrypsin digestion, which removed a substantial amount of non-fibrillar proteins, the collagen banding pattern became visible (**D**).

We then used amino acid analysis to confirm that, after chymotrypsin digestion, the ECM was substantially enriched in collagen. The amino acid composition of collagen differs significantly from that of other proteins because every third amino acid in the collagen triple helix is glycine (Gly) [22,27]. Proline (Pro) and alanine (Ala) are also unusually abundant in collagen [27]. Hydroxylated amino acids—hydroxyproline (Hyp) and hydroxylysine (Hyl)—are characteristic of collagen and exceptionally rare in non-collagenous proteins [27]. Therefore, after chymotrypsin treatment, an increase in the relative content of glycine, proline, alanine, hydroxyproline, and hydroxylysine were expected in the ECM, alongside a decrease in leucine (Leu), isoleucine (Ile), phenylalanine, and tyrosine, as these amino acids are less abundant in collagen compared to non-collagenous ECM proteins [27]. Indeed, amino acid analysis showed that with each cleaning step, the ECM became progressively enriched in collagen with the relative content of collagen-specific amino acids gradually approaching their expected values (Table 2). The slight discrepancies in amino acid composition of the chymotrypsin-treated samples compared to purified collagen type I could be attributed to the presence of different types of collagen in the ECM and the different species each ECM was derived from. Thus, amino acid analysis correlated well with fluorescence measurements, demonstrating that chymotrypsin successfully digested all proteins except collagen.

**Table 2.** The molar percentage (mol%) of amino acids in FSOB and BVSMC ECM.

| Amino<br>Acid<br>residues | Collagen<br>type I<br>from<br>Achilles<br>tendon | BVSMC |  |  | FSOB |  |  |
| --- | --- | --- | --- | --- | --- | --- | --- |
|  |  | Chymotrypsin | Detergent | Freeze-thaw | Chymotrypsin | Detergent | Freeze-thaw |
| *High relative abundance in collagen* |  |  |  |  |  |  |  |
| Ala | **8.27** | **7.94** | 5.63 | 5.59 | **7.79** | 6.56 | 5.41 |
| Gly | **22.00** | **20.25** | 9.90 | 9.89 | **19.41** | 13.39 | 7.81 |
| Pro | **12.22** | **12.24** | 7.52 | 7.62 | **11.42** | 8.94 | 6.43 |
| Hyp | **11.16** | **11.94** | 4.15 | 4.21 | **11.09** | 7.17 | 2.54 |
| Hyl | **1.00** | **1.61** | 0.44 | 0.39 | **1.73** | 0.77 | 0.37 |
| *Low relative abundance in collagen* |  |  |  |  |  |  |  |
| Leu | **3.42** | **3.38** | 7.01 | 6.99 | **3.74** | 5.71 | 7.88 |
| Ile | **1.58** | **1.68** | 3.67 | 3.82 | **1.79** | 2.98 | 4.20 |
| Phe | **2.35** | **2.27** | 3.69 | 4.15 | **2.53** | 3.30 | 4.36 |
| Tyr | **0.79** | **0.70** | 3.31 | 3.46 | **1.06** | 2.36 | 3.84 |
| *Other* |  |  |  |  |  |  |  |
| Arg | **8.74** | **8.96** | 8.83 | 8.34 | **8.87** | 9.17 | 8.84 |
| Asp | **5.88** | **5.90** | 8.65 | 8.82 | **6.33** | 8.04 | 9.45 |
| Glu | **10.66** | **10.76** | 13.45 | 12.81 | **10.83** | 12.11 | 12.49 |
| His | **0.77** | **0.83** | 2.19 | 2.18 | **0.88** | 1.58 | 2.26 |
| Lys | **3.43** | **2.93** | 5.94 | 5.79 | **3.33** | 5.12 | 7.30 |
| Met | **0.09** | **0.94** | 1.97 | 1.89 | **0.97** | 1.61 | 1.91 |
| Ser | **3.11** | **3.07** | 4.39 | 4.52 | **3.03** | 3.76 | 4.68 |
| Thr | **1.91** | **2.00** | 4.39 | 4.50 | **2.34** | 3.45 | 4.63 |
| Val | **2.61** | **2.60** | 4.88 | 5.05 | **2.86** | 3.98 | 5.61 |

Mol% was measured by amino acid analysis as described in Methods. ECM was progressively cleaned by freeze-thaw cell lysis, detergent disruption of lipid-lipid/protein interactions, and chymotrypsin digestion of non-collagenous proteins (see in Methods). Due to hydrolysis, amino acids aspartate and asparagine (Asn) were combined as Asn, and glutamate and glutamine (Gln) as Glu. Data were normalized against L-norleucine, which was used as an internal standard. Collagen type I, purified from bovine Achilles tendon, was obtained from Sigma.

Transmission electron microscopy (TEM) can visually demonstrate the presence of intact collagen fibrils. In TEM images, collagen fibrils appear as long (typically longer than the field of view), relatively straight (i.e., not kinked) fibril structures and exhibit a distinctive banding pattern [28]. These unique features of collagen fibrils enable their accurate recognition in TEM images. Images of the *in vitro* ECM after freeze-thaw cell lysis and detergent washes showed that a substantial amount of non-fibrillar protein remained in the sample (Fig. 1C). Collagen fibrils were observed, but the banding pattern was not visible, as it was obscured by non-fibrillar proteins attached to the collagen fibrils. After majority of non-collagenous proteins were removed by chymotrypsin treatment (Fig. 1D), collagen fibrils could be easily identified not only by their straightness and length, but most importantly, by their unique banding pattern.

Next, we used solid-state NMR spectroscopy (ssNMR) to demonstrate that the bulk collagen triple helix structure was indeed preserved after our clean-up procedure. ^13^C ssNMR provides structural information about the bulk of the sample, as different amino acid residues produce ^13^C signals at distinct resonance frequencies, and are sensitive to secondary protein structures [29], with triple-helical collagen proteins exhibiting distinctive signal patterns due to their high glycine, proline, and hydroxyproline content and their unique helical packing [30].

After the chymotrypsin step the overall ^13^C ssNMR spectrum of ECM-extracted collagen was indistinguishable from commercially-available purified collagen type I (Fig. 2A and S1). Firstly, after each step of the clean-up procedure, the ^13^Cα and ^13^C’ NMR signals from Gly and Pro, along with the ^13^Cγ and ^13^Cβ signals of Hyp, became progressively more prominent, with every ^13^C chemical shift for Pro, Hyp, Ala, and Gly becoming increasingly typical of the collagen triple helix [31]. Secondly, and most importantly, we compared ssNMR ^13^C spectra of ECM-extracted collagen with commercial collagen type I, which had been partially digested, i.e. denatured, by collagenases (Fig. 2B). In the denatured collagen spectrum, significant signal shifts of Gly ^13^Cα and Pro ^13^Cα, as well as overall peaks’ broadening, were observed, revealing the spectral signature of the denatured collagen triple helix (Fig. 2B). However, no obvious changes in ^13^C chemical shifts or relative signal intensities were observed in the ssNMR spectrum of collagen, purified by chymotrypsin (Fig. 2B), indicating that the bulk triple helical structure was indeed preserved and no significant denaturation occurred during out chymotrypsin treatment.

**Fig. 2.**
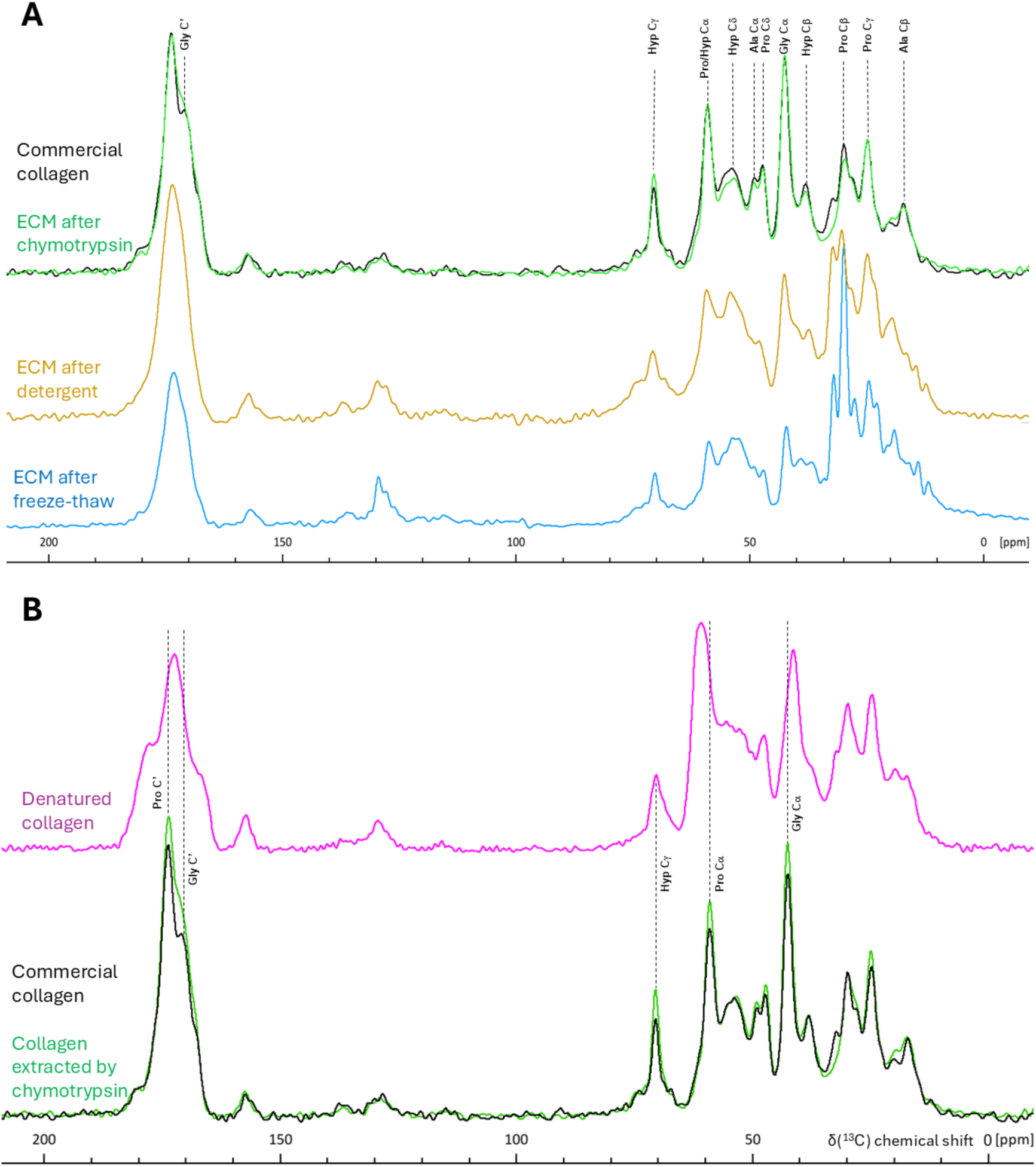
¹³C CP/MAS ssNMR spectra of unlabelled/natural abundance BVSMC ECM following collagen isolation steps. **A.** Collagen isolation steps included either freeze-thaw cell lysis or Triton X-100 cell lysis (“detergent”), followed by chymotrypsin digestion. The control spectrum was represented by commercially available bovine collagen type I. After the chymotrypsin step, collagen from the ECM was clean enough to correlate well with that of commercial collagen. **B.** After partial digestion of commercial collagen with collagenase, the resulting denatured collagen shows shifted proline and glycine signals compared to both commercial collagen and ECM collagen extracted *via* chymotrypsin. All samples were freeze-dried and spectra were recorded on a 400 MHz spectrometer at room temperature using a 10 kHz MAS rate, and the following number of scans: 20k (freeze-thaw), 10k (denatured collagen), 8k (both detergent and chymotrypsin), and 256 (commercial collagen).

**SUPPLEMENTARY FIGURE. Fig. S1.**
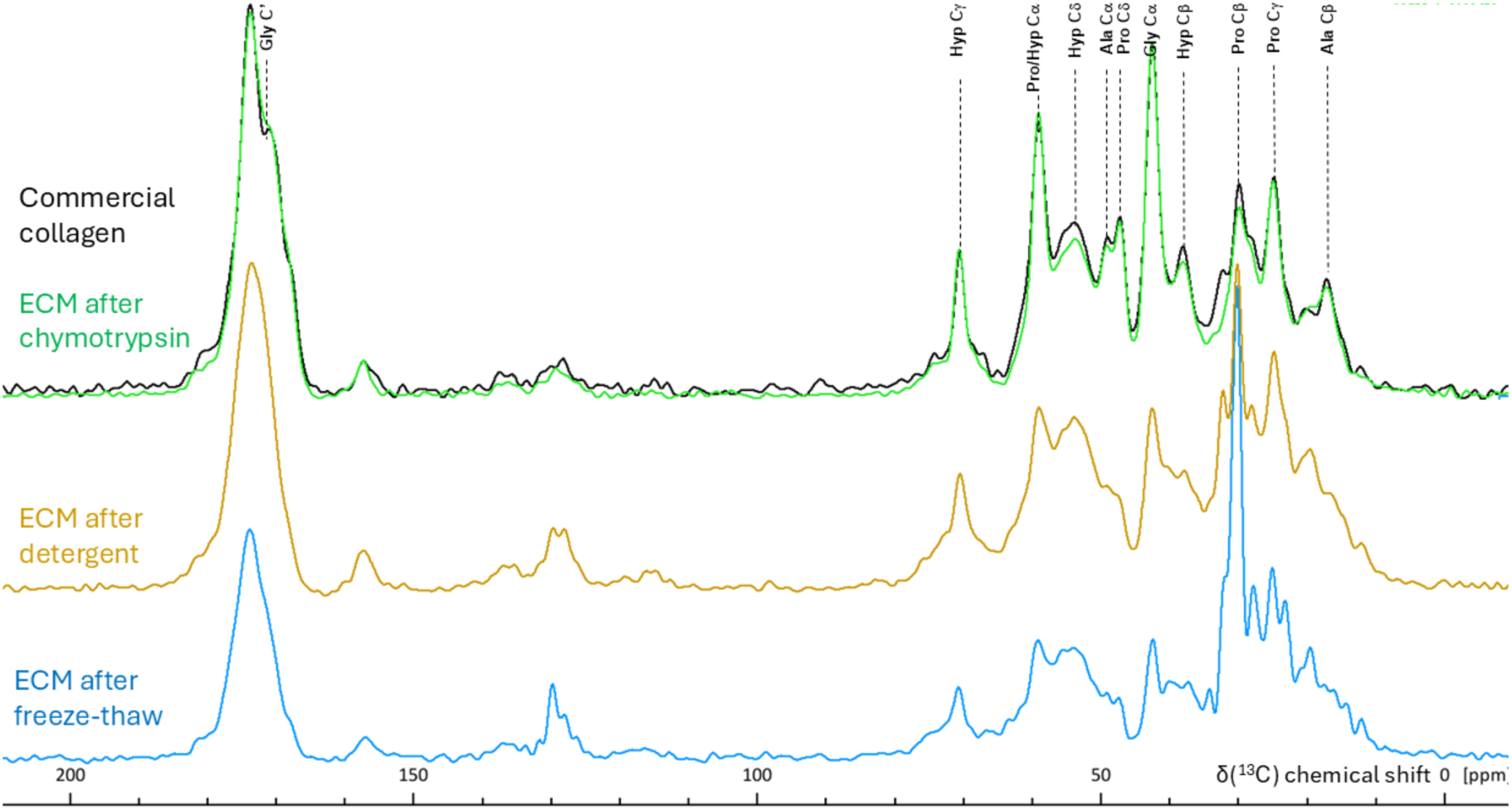
¹³C CP/MAS ssNMR spectra of unlabelled/natural abundance FSOB ECM following collagen isolation steps. Collagen isolation steps included either freeze-thaw cell lysis or Triton X-100 cell lysis (“detergent”), followed by chymotrypsin digestion. The control spectrum was represented by commercially available bovine collagen type I. Only after the chymotrypsin digestion step, collagen from the ECM was clean enough to correlate well with that of commercial collagen. Samples were freeze-dried and spectra were recorded on a 400 MHz spectrometer at room temperature using a 10 kHz MAS and the following number of scans: 32k (chymotrypsin), 8k (both freeze-thaw and detergent) and 256 (commercial collagen).

To summarize, collagen with a preserved triple helix structure in well-defined fibrils was the bulk protein remaining in the FSOB and BVSMC ECMs after purification, with the crucial step being the digestion of non-collagenous proteins by chymotrypsin. Protein autofluorescence enabled us to determine the optimal digestion duration, while amino acid analysis confirmed that after digestion the ECM had amino acid ratios typical of collagen. NMR spectroscopy confirmed the bulk preservation of the triple helix structure, while TEM visualized the depletion of non-collagenous proteins, with collagen fibrils retaining their characteristic banding.

### Decellularization to obtain native ECM

We then developed a decellularization procedure tailored for studies in which a native ECM is essential [6,7,12,13]. Our goal was to preserve the ECM’s composition and protein intertwining as closely as possible to the pre-decellularization state. To achieve this, we designed a strategy that dislodges unruptured living cells from the ECM using cytoskeleton-targeting drugs, thereby avoiding contamination with intracellular material. This approach eliminates the need for the relatively harsh treatments typically used to remove cell debris following lysis—treatments that often result in significant loss of soluble ECM proteins and thus compromise the retention of native protein network in ECM [7].

Initially, we tested vinblastine, which is known to depolymerize microtubules [32]. Microtubules interact with both actin and keratin filaments [33,34], which in turn support adhesion structures such as focal adhesions and hemidesmosomes. Therefore, microtubule collapse could potentially lead to the disassembly of all adhesion complexes. Indeed, addition of vinblastine induced filopodia retraction and cell rounding in three different cell cultures (Fig. 3).

**Fig. 3.**
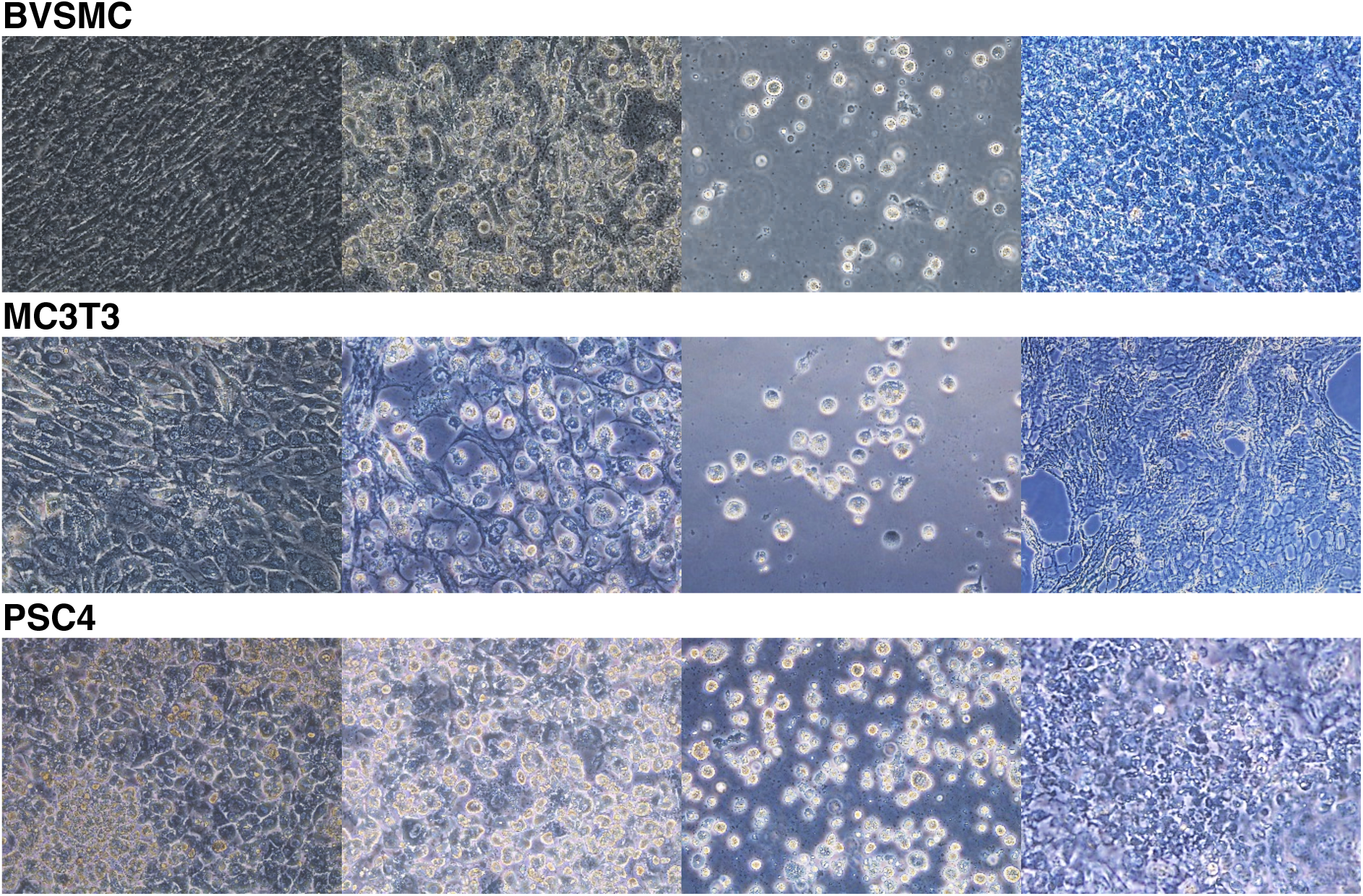
Brightfield microscopic observations of the ECM decellularization process using cytoskeleton-targeting drugs. After reaching confluency, cell cultures were maintained for 3–4 weeks in the presence of ascorbic acid to promote collagen production. Cells were then treated with 80 mM vinblastine sulphate (PSC4); 360 mM vinblastine sulphate and 35 mM sodium orthovanadate (BVSMC); and 120 µM vinblastine sulphate, 35mM sodium orthovanadate, and 62 µM latrunculin B (MC3T3). Loss of cell– ECM adhesion and subsequent cell rounding were observed as birefringence at the edges of cell bodies. Detached cells appeared rounded, while the cleaned ECM appeared as an acellular sheet (BVSMC and PSC4) or as a “fishing net”-like tissue with numerous holes (MC3T3), corresponding to areas previously occupied by cells. Each field of view is 475 µm × 624 µm. Representative images were captured using a Leica DMi1 brightfield microscope.

However, treatment with vinblastine alone was insufficient for complete removal of cells (Fig. 4), as many rounded cells remained attached to the ECM. We hypothesized that while the actin cytoskeleton in the filopodia was sufficiently depolymerized—evidenced by filopodia retraction—the depolymerization of keratin filaments in hemidesmosomes was not achieved, causing rounded cells to either stay attached to the ECM or be dislodged together with ECM fragments. To assist keratin depolymerization, we decided to add sodium orthovanadate, a disruptor of the keratin filament network [35]. Co-application with vinblastine resulted in effective ECM clean-up in both PSC4 and BVSMC cultures. Compared to vinblastine alone the number of detached cells increased by 60% (Fig. 4, BVSMC). Decellularized ECMs contained many structures that could be mistaken for residual cells, especially in PSC4. However, Hoechst staining revealed no nuclei in either ECM following the most effective cell-removal treatments (Fig. 4, BVSMC and PSC4), making the presence of cells unlikely, as nuclei—whether healthy, apoptotic, or damaged—typically produce a strong Hoechst signal. Increasing the laser excitation intensity threefold revealed numerous oval or angular structures, consistent with weak autofluorescence from amino acids such as tyrosine and tryptophan and their derivatives. While these could theoretically represent apoptotic bodies retained within the ECM, the complete absence of Hoechst-positive nuclear debris makes this interpretation unlikely. Altogether, the data support the conclusion that these weakly fluorescent structures likely consist of ECM rather than cellular proteins.

**Fig. 4.**
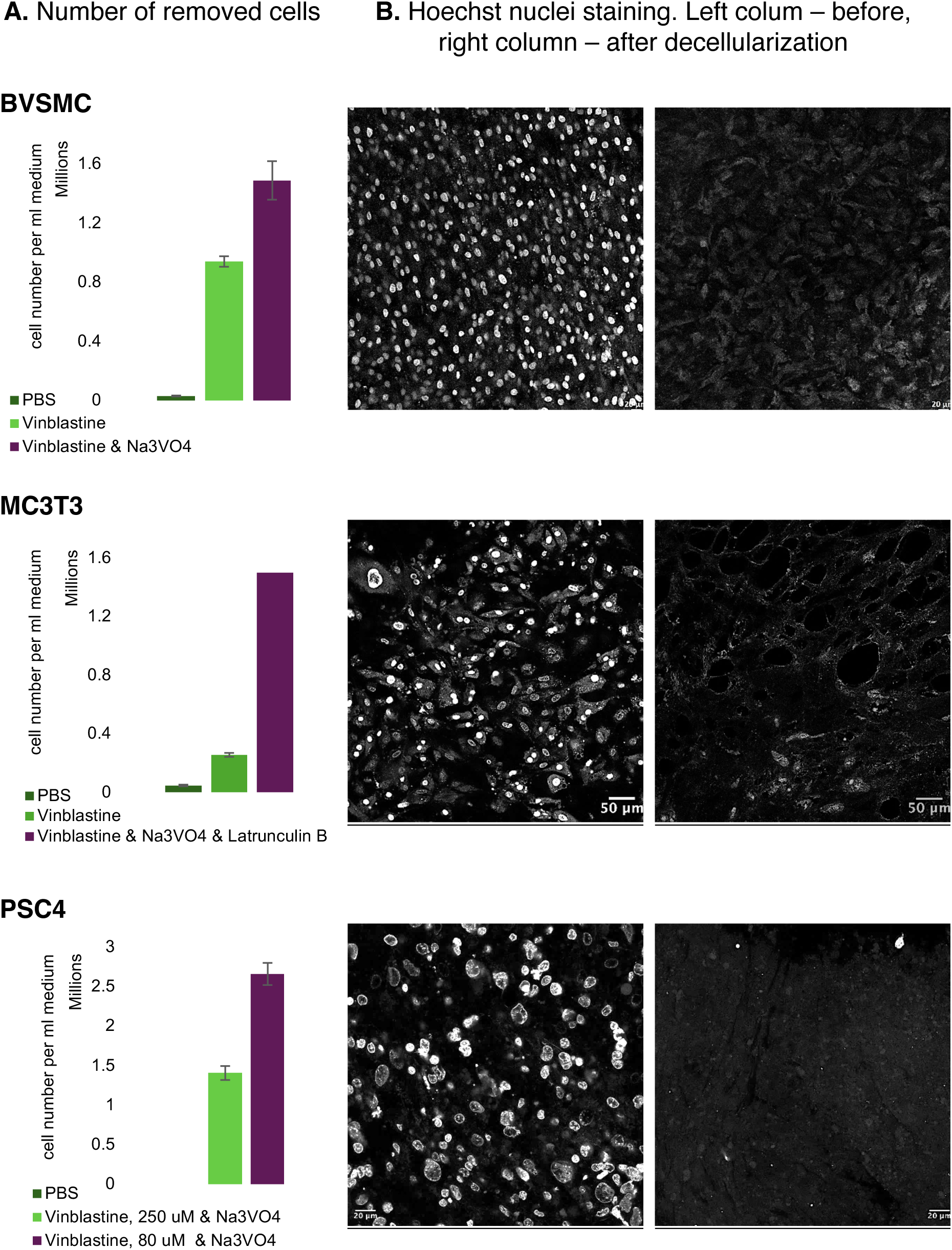
Validation of ECM decellularization. **A.** Different cell cultures required different combinations and concentrations of cytoskeletal-targeting drugs for maximal removal of cells from the ECM. Bar charts represent the average and standard error of two to four independent experiments. **B.** Effective decelllarization was confirmed by Hoechst 33342 staining of nuclei in the ECM before (left column) and after (right column) treatment with cytoskeleton-targeted drugs. The most effective treatments were as follows: 80 mM vinblastine sulphate (PSC4); 360 mM vinblastine sulphate and 35 mM sodium orthovanadate (BVSMC); and 120 µM vinblastine sulphate, 35 mM sodium orthovanadate, and 62 µM latrunculin B (MC3T3). Representative images were captured using a Leica STELLARIS 5 confocal microscope.

It is important to note that vinblastine concentration played a critical role in BVSM and PSC4 decellularization. When testing for the optimal dose, we found that in the case of PSC4, a concentration of 80 µM resulted in greater cell detachment than 250 µM (Fig. 4, PSC4), most likely because the higher dose was cytotoxic, leading to cell death rather than physiological microtubule disruption. In contrast, for BVSMCs, lower vinblastine concentrations were ineffective at inducing cell rounding (data not shown), and a higher concentration (360 µM) was required to achieve microtubule depolymerization. This highlights that each cell culture has a unique tubulin sensitivity to co-crystallisation with vinblastine (see more in Discussion), necessitating careful optimization in each case. We also briefly tested various concentrations of sodium orthovanadate (data not sown) and found 35 mM to be optimal for all cultures. This is possibly because the intracellular orthophosphate concentration is controlled by its level in the medium, thereby also determining the required concentration of its competing ion, orthovanadate.

However, in the case of the MC3T3 culture, the combination of vinblastine and orthovanadate did not result in well-decellularized ECM, and many cells did not round properly and remained attached. We hypothesized that in this case, microtubule collapse alone was insufficient to fully depolymerize actin filaments. Therefore, we tested the actin-disrupting agent latrunculin B [35]. Also, to maintain cell viability during treatment with three cytoskeleton drugs, we chose not to apply all three agents simultaneously, but sequentially. We first applied vinblastine, followed by latrunculin B, and then treated the cells with sodium orthovanadate. Additionally, we shortened the duration of each treatment and allowed the cells to rest overnight in fresh medium. This modified protocol proved effective: under the brightfield microscope, the cells appeared fully rounded (Fig. 3, MC3T3). The purified MC3T3 ECM exhibited a markedly different structure compared to that observed for BSMC and PSC4. In these two cell cultures, the ECM formed a continuous layer with no interruptions beneath the cells (Fig. 3). In contrast, the ECM produced by MC3T3 cells appeared as a mesh of fibres surrounding, but not underlying, the cells (Fig. 3). As a result, the decellularized MC3T3 ECM resembled a “fishing net” (Fig. 3, MC3T3), with numerous holes corresponding to areas previously occupied by cells. In the confocal images, these holes were also clearly visible as black circular regions completely lacking the weak autofluorescence of ECM proteins (Fig. 4, MC3T3).

Importantly, following decellularization, the BVSMC and MC3T3 ECMs remained structurally intact and could be handled as single pieces of tissue. In contrast, the PSC4 ECM, which lacks fibrillar collagen and was not expected to remain fully cohesive, fragmented into several pieces. However, these fragments could still be handled with tweezers and did not disintegrate. This suggests that within the matrix, ECM proteins were still held together by weak intermolecular interactions. Overall, these findings indicate that the protein network was preserved across all three ECMs.

To confirm that our new decellularization procedure preserved non-collagenous proteins within the ECM, we employed ssNMR—this time applying it to frozen ECM rather than freeze-dried collagen samples, in order to better preserve networks composed of different proteins and stabilized by weak hydrophobic and hydrophilic interactions. Performing ssNMR on fully hydrated tissue is challenging, as the high salt content limits the radiofrequency power that can be applied and leads to sample heating [37]. Additionally, the magic-angle spinning (MAS) required for ssNMR can distort the structure of softer ECM components. To overcome these limitations, we froze the ECM samples, which allowed us to record cross-polarization CP/MAS ¹³C ssNMR spectra with good signal-to-noise ratios.

The ^13^C CP/MAS ssNMR spectrum of BVSMC ECM was expected to be dominated by fibrillar collagen (such as collagen type I and III), as our preliminary proteomics showed these cells produce it in substantial amounts (data not shown). Indeed, the decellularized ECM spectra contained all the characteristic features of fibrillar collagen (Fig. 5A). First, all hydroxyproline signals (Hyp ^13^Cα, ^13^Cβ, ^13^Cγ, and ^13^Cδ) were well-defined in the decellularized ECM spectra, confirming the presence of collagen. Notably, the Hyp ^13^Cβ and ^13^Cδ peaks were better resolved in the frozen ECM than in the completely purified dry collagen control (compare Fig. 5A and Fig. 2A), evidencing greater molecular organization of collagen in the decellularized ECM sample. This is consistent with the retention of essential weak hydrophilic and hydrophobic interactions between collagen and other ECM proteins, for example, proteoglycans, as previously demonstrated [38]. In the ^13^C ssNMR spectrum of non-decellularized ECM, the relative intensity of the Hyp signals was diminished, and these signals were completely absent in the spectrum of cells, confirming the effective removal of cells without dislodging the ECM (Fig. 5A). Second, the Gly ^13^Cα was the most prominent signal in the decellularized ECM (Fig. 5A), but was absent in the cell spectrum, consistent with the low glycine abundance in intracellular proteins. Third, ^13^C signals from proline and alanine, were well-defined in the decellularized ECM spectrum (Fig. 5A), consistent with the high relative abundance of these residues in ECM proteins, and collagens in particular. Fourth, the ^13^C chemical shifts of Gly, Pro, and Hyp in the decellularized BVSMC ECM spectrum (Fig. 5B) were typical of those for the collagen triple helix [31].

**Figure 5.**
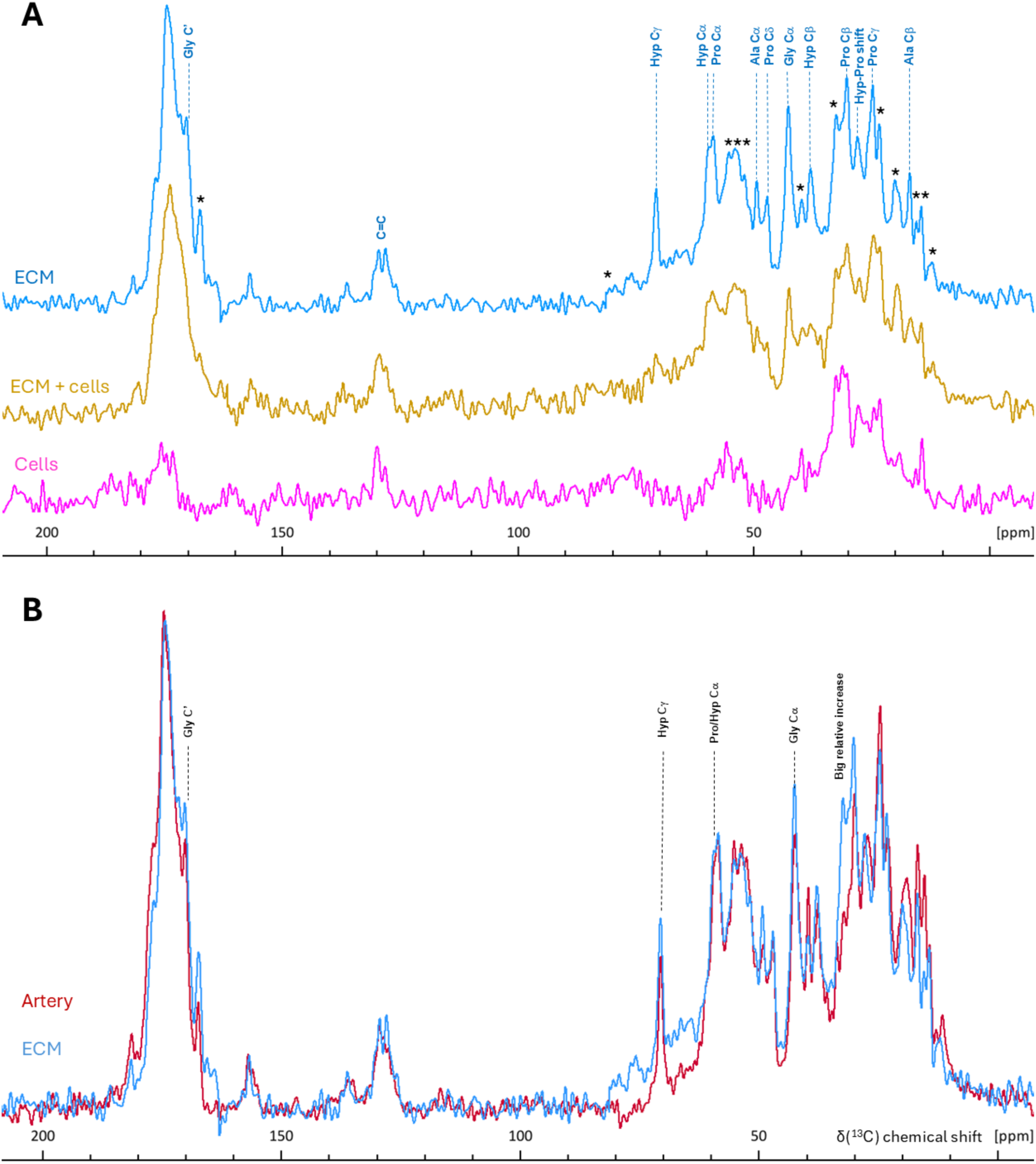
¹³C CP/MAS ssNMR spectra of unlabelled detached cells, non-decellularized ECM, and decellularized ECM from BVSMCs, compared with bovine tail artery. **A.** Cells showed no collagen-specific spectral features, whereas these features were prominent in the decellularized ECM. Asterisk (**\***) marks the presence of non-collagen signal in the spectrum of the decellularized ECM. **B.** The spectrum of decellularized ECM closely matched that of the bovine tail artery. Spectra were acquired on a 400 MHz spectrometer using MAS rate of 11 kHz at −8°C (corrected for friction heating) and the following number of scans: 46k (cells) and 16k (ECM or ECM + cells), and 24k (artery).

Importantly, the ^13^C ssNMR spectrum of the decellularized BVSMC ECM contained features that were absent in pure collagen type I, such as signals at 12.2, 14.4, 15.4, 20, 23.3, 32.4, 39.7, 51.8, 53.9, 55.2, 79.3 and 167.5 ppm (Fig. 5A). By using our decellularization procedure, which dislodges whole cells or, towards the end, apoptotic bodies [39], we can be certain that the observed signals originate from ECM proteins, not cellular debris. Our preliminary proteomic and immunocytochemical studies demonstrated (data not shown) presence of many ECM proteins, such as collagen I, V, VI, XI, and XII, elastin, fibulins, laminins, fibronectin, decorin, perlecan, lumican, biglycan and tenascin C in BVSMC cultures. Hence, in carefully decellularized BVSM ECM, there is potentially many proteins that could generate ¹³C NMR signals with chemical shifts distinct from those of the collagen triple helix. Assigning every unknown feature in the spectrum could take several years of research. However, a simpler approach to confirm that the chemical shifts listed above indeed represent ECM proteins was to compare them with those from *ex vivo* tissue, where the ¹³C ssNMR signal from ECM overwhelmingly dominates over cellular components. A good example of such tissue is the artery wall, where the ECM produced by the same VSMC is the major constituent. Serial block-face scanning electron microscopy 3D volume images have shown that the medial lamellar unit (MLU)—the fundamental structural and functional unit of the aorta— was comprised, by volume, of 47% collagen, 29% elastin, and only 24% VSMCs [40]. Comparison of the ^13^C ssNMR spectral features of the ox tail artery with the decellularized BVSMC ECM showed that the spectra were nearly identical with two small discrepancies: a broad signal in the region at 60-80 ppm, and a very high intensity of an aliphatic signal at 32.4 ppm (Fig. 5B). However, neither of these features were observed in the dislodged cells (Fig. 5A), suggesting that they represented differences between *in vitro*-grown ECM and the *ex vivo* matrix of the ox tail artery. Thus, the unassigned, non-collagen spectral features in the ^13^C ssNMR spectrum of decellularized ECM (Fig. 5A) indeed corresponded to non-collagenous ECM proteins. Furthermore, the decellularized ECM appears to be a good *in vitro* model of the artery MLU in terms of the ECM protein composition and structure.

The ¹³C ssNMR spectrum of the decellularized, non-activated PSC4 ECM was dominated by features arising from non-collagenous proteins with some collagen features— particularly Pro/Hyp ^13^Cα and Hyp ^13^Cγ—present but of relatively weak intensity (Fig. 6). Notably, the characteristic Gly ^13^Cα signal at 42.5 ppm, typically associated with fibrillar collagen, was almost absent, suggesting that Pro/Hyp signals come from flexible, non-fibrillar collagens such as, for example, types IV and VI. We expected to observe such a “collagen-poor” spectrum because PSC4 was cultured without activation factors and, under these conditions, did not show any staining for procollagen III or collagen I [41]. A variety of structural proteins in PSC ECM would include tenascin, fibronectin, laminins, fibrillins, and ECM1, as well as non-fibrillar collagens (types IV, VI, XVI, and XXIII) and fibrillar collagens (types I, II, III, and V), as shown by proteomics [42]. All of these proteins, except fibrillar collagens, can potentially generate ¹³C ssNMR signals in the decellularized, non-activated PSC4 ECM.

**Fig. 6.**
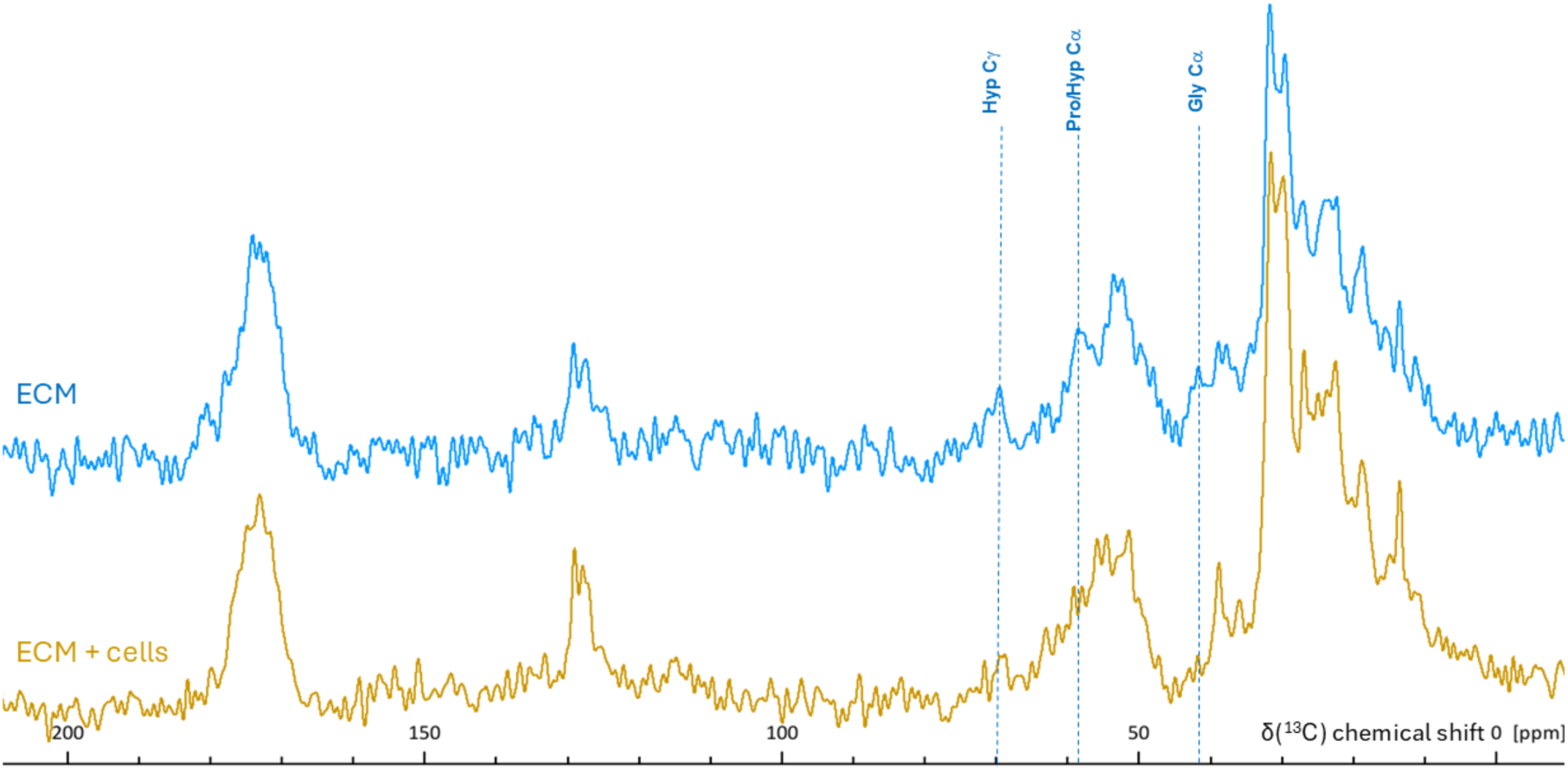
¹³C CP/MAS ssNMR spectra of unlabelled ECM from PSC4 culture. The non-decellularized ECM showed no collagen-specific spectral features, whereas collagen specific signals (Hyp Cα, Hyp Cγ, Pro Cα, and Gly Cα) became detectable after decellularization. However, non-collagen features continued to dominate the spectrum. Spectra were acquired using a 400 MHz spectrometer with MAS rate of 12 kHz, at –10 °C (corrected for frictional heating), and 10k scans.

Thus, we can conclude that our novel decellularization procedure, which dislodges whole living cells using cytoskeleton-targeting drugs, yields a clean ECM while preserving its native complexity—whether it is collagen-rich or collagen-poor.

## DISCUSSION

Many extracellular matrix-related diseases, such as atherosclerosis, arteriosclerosis, fibrosis, cancer spread, and metastasis, require molecular cell biology studies with cells in their native or near-native ECM environment. Novel cell therapies like stem cell transplantation and tumour-infiltrating lymphocyte (TIL) and CAR T-cell therapies, require testing of their effectiveness within native extracellular matrix settings [43–46]. For instance, studies have shown that increased rigidity of the ECM can impair the function of T cells [46], highlighting the necessity of the ECM in the design of therapies.

Though ECM is never composed solely of collagen (see Introduction), it is often a major protein that determines cell anchoring, mobility and fate, as collagen contributed to the evolution of multicellular animals [47]. It is no wonder that cells have numerous receptors which can bind directly to collagen or its sugar chains. Collagen sensors – tyrosine receptor kinases discoidin domain receptors 1 and 2 (DDR1 and DDR2) - are present in many tissues [48]. The largest family of collagen adhesion receptors consists of αI domain-containing integrins [47], which are present in focal adhesion sites in nearly every vertebrate cell. Collagen binding to annexin V regulates mineralisation in chondrocytes [49]. Collagen binding to platelet receptors glycoprotein Vi (GPVI) and CD36 is an important stage in thrombosis [50] The high avidity collagen receptor - inhibitory leukocyte-associated immunoglobulin-like receptor-1 (LAIR-1) - sets threshold for activation of natural killer cells, effector T cells, B cells and dendritic cells, and also may down-regulate responses directed against the tumour by various effector cells [51]. Cell surface heparan sulphate proteoglycans, such as syndecan-1 and −2, and glypican-1, bind to sugar residues on collagens [52] providing sugar-specific type of binding.

Therefore, seeding cells onto collagen matrices is used in numerous research studies: a basic search in Science Citation Index Expanded yields over 63,000 papers containing phrase “cells in collagen matrix” [53] highlighting the importance of the subject. However, commercially-available reconstituted collagens (mainly bovine and rat) cannot be considered native because proteolysis damages the fibril structure, and acid hydrolysis breaks glycosidic bonds in post-translational modifications. Moreover, in most cases, such collagen is neither species- nor tissue-specific. In this paper, we report a new method for collagen purification from *in vitro*-grown ECM, which could potentially be applied to any type of cell capable of producing collagen *in vitro*. This method expands the range of species- and tissue-specific collagens available for research, making it especially important for human-related studies and drug discovery. We show that this method preserves the collagen fibril structure and is unlikely to damage its post-translational modifications. When produced from patient-derived cells, such collagen— being a syngeneic biological material—can offer the highest possible biocompatibility [54].

For applications where native ECM is essential [43–46], we developed a novel decellularization procedure, exploiting a physiological approach to cell detachment from the ECM, yielding a clean native ECM. Methods that use cell lysis result in DNA and other nuclear material adventitiously adhered to the ECM. We have previously demonstrated that it is impossible to completely remove DNA and nuclei from the ECM using conventional methods [10]. Furthermore, methods which involve cell lysis exposes the ECM to aggressive intracellular enzymes (e.g., proteases, phosphatases, etc.), which leads to ECM modification and also involve aggressive chemicals that can break ester and glycosidic covalent bonds in the ECM, leading to ECM degradation [1,2].

To drive the detachment of live cells from the ECM, we chose a strategy based on disrupting the cytoskeletal architecture, leading to the loss of cell adhesion. In general, cell adhesion is mediated by focal adhesions (FAs) and podosomes supported by actin filaments [55], as well as hemidesmosomes anchored by keratin intermediate filaments [56]. Both types of filaments engage in cross-talk with microtubules through molecular components that provide a physical crosslink [33,34]. Hence, our initial approach was to trigger microtubule depolymerization and, consequently, collapse the actin- and keratin-based adhesive structures.

Although the majority of studies on tubulin focus on the inhibition of microtubule elongation, an interesting study on vinblastine-treated cancer cell lines demonstrated that, at concentrations well above cancer-related IC50, it was possible to induce fragmentation of already assembled microtubules [57]. Furthermore, in 1968, it was shown that vinblastine, at isodesmic concentrations, induced the formation of paracrystalline tubulin aggregates in oocytes, fibroblasts, and leukocytes [32]. A similar effect was observed with extracted tubulin: vinblastine at stoichiometric concentrations induced paracrystals similar to those formed in oocytes [58]. In our study, the range of concentrations of vinblastine (80-360 μM, Fig. 1) required for cell rounding correlated with the mode of action of vinblastine reported back in 1968, where it was shown to induce the formation of paracrystalline tubulin aggregates.

However, vinblastine alone did not dislodge all cells from the matrix, and forcing them out by vigorous shaking resulted in numerous ECM ruptures. These observations led us to hypothesize that cell-ECM junctions—specifically keratin component of hemidesmosomes [59] —was not fully disassembled during vinblastine treatment. The balance between the different organizational states of keratins (filamentous, granular, soluble) depends on the state of cellular phosphorylation. Therefore, tyrosine kinase inhibitor sodium orthovanadate and the serine/threonine phosphatase inhibitor okadaic acid were considered, as both are known to disrupt the keratin filaments and lead to reversible formation of granular aggregates [60–62]. For ECM decellularization, we decided to use sodium orthovanadate, because upon this treatment, plectin was shown to colocalize with keratin granules, suggesting a better chance of hemidesmosome disassembly [61]. Indeed, the combination of sodium orthovanadate with vinblastine resulted in decellularized ECM in the case of BVSMC and PSC4 cultures (Fig. 3 and 4).

In the case of MC3T3 cells, the additional application of the actin-targeting drug latrunculin B [63] was found to be essential. This may be due to its dual mechanism of action: it not only prevents actin polymerization but also disrupts focal adhesion formation by blocking the translocation of FAK and paxillin to adhesion sites [63]. It is possible that microtubule collapse induced by vinblastine was insufficient to destabilize the actin cytoskeleton, or that actin structures were rapidly reassembled in this particular cell line.

The most widely used *in vitro* model of native ECM is Matrigel®, an extract from Engelbreth–Holm–Swarm mouse sarcoma. It has been utilized in over 12,000 publications [64] and has provided scientists with the opportunity to study embryonic, normal, stem, and malignant cells in a more natural 3D environment compared to growing them on a plastic surface in liquid growth media. However, the composition of Matrigel® (60% laminins LAMB1, LAMA1, and LAMC1; 20% nidogen NID1; and 20% fibrinogens FGG, FGB, and FGA [65], does not reflect the structural complexity of ECMs, nor can it reflect ECM changes in matrix-related diseases and therapies. For instance, it has been shown that gastrointestinal organoids grow and transplant better when cultured in hydrogels derived from decellularized gastrointestinal tissues compared to Matrigel® [65]. This is understandable because decellularized gastrointestinal tissues mainly contained collagens and proteoglycans, which Matrigel® lacked.

Our method enables the production of native ECM—either collagen-rich or collagen-poor—from *in vitro*-grown cells. This significantly expands the range of native matrices available for 3D studies. The resulting ECM can be dried, powdered, and reconstituted into a gel, or used as a sheet for repopulation with cells of interest. It also offers potential for tissue engineering applications, where it can be combined with synthetic polymers to address the limitations of polymer-only biocompatibility and the insufficient mechanical strength of ECM-only constructs [66].

## CONCLUSIONS

In this study, two novel methods for isolating native extracellular proteins were developed. Collagen with a preserved triple-helical structure and well-defined fibrillar organization was successfully purified from FSOB and BVSMC cell cultures, with the digestion of non-collagenous proteins by chymotrypsin identified as a critical step. In parallel, intact ECM with preserved protein complexity was isolated from BVSMC, MC3T3, and PSC4 cell cultures using a newly developed decellularization strategy that removes entire living cells via the application of cytoskeleton-targeting agents, including vinblastine, sodium orthovanadate and latrunculin B.

## AVAILABILITY OF DATA

Raw ssNMR experimental data files are available in the Cambridge Research Repository, Apollo, with the identifier: https://doi.org/10.17863/CAM.120792

Other raw data for this article is available upon request from the corresponding author.

## ACKNOWLEDGEMENTS

The authors would like to thank Dr Peter Sharratt for assistance with amino acid analysis, Dr Karin Mueller for assistance with TEM, Dr Sneha Bansode for providing collagenase-denatured collagen, Dr Rakesh Rajan for isolation of FSOB, Prof. Catherine Shanahan (King’s College London) for the donation of BVSMCs, and Dr David Reid for helpful discussions.

## AUTHOR CONTRIBUTIONS

Uliana Bashtanova: conceptualization, methodology, data acquisition & analysis, and writing - original manuscript. Rui Li, Ieva Goldberga, Kathryn Gerl, Kristen Burgess, and Annika Wegner-Repke: data acquisition & analysis, writing - review & editing. Melinda Duer: funding acquisition, project administration, methodology, writing – review & editing.

## CONFLICT OF INTREST

The authors declare no conflict of interest

